# Common and distinct neural correlates of music and food-induced pleasure: a coordinate-based meta-analysis of neuroimaging studies

**DOI:** 10.1101/2020.08.14.250894

**Authors:** Ernest Mas-Herrero, Larissa Maini, Guillaume Sescousse, Robert J. Zatorre

## Abstract

Neuroimaging studies have shown that, despite the abstractness of music, it may mimic biologically rewarding stimuli (e.g. food) in its ability to engage the brain’s reward circuity. However, due to the lack of research comparing music and other types of reward, it is unclear to what extent the recruitment of reward-related structures overlaps among domains. To achieve this goal, we performed a coordinate-based meta-analysis of 38 neuroimaging studies (703 subjects) comparing the brain responses specifically to music and food-induced pleasure. Both engaged a common set of brain regions including the ventromedial prefrontal cortex, ventral striatum, and insula. Yet, comparative analyses indicated a partial dissociation in the engagement of the reward circuitry as a function of the type of reward, as well as additional reward type-specific activations in brain regions related to perception, sensory processing, and learning. These results support the idea that hedonic reactions rely on the engagement of a common reward network, yet through specific routes of access depending on the modality and nature of the reward.

## INTRODUCTION

Humans have the capacity to feel and experience pleasure from a large variety of stimuli and activities: from primary rewards that satisfy basic biological drives, such as food, to aesthetic experiences, such as listening to music. All rewarding events and experiences, independently of their type – from primary to aesthetic – are processed by a common set of brain regions that constitute the well-known reward circuit, including the ventral tegmental area, the ventral striatum, the insula, and the ventromedial prefrontal cortex, among others (Bartra et al., 2013; Blood and Zatorre, 2001; Sescousse et al., 2013). This circuitry, through a complex interplay between dopaminergic and opioid pathways, is involved in many different aspects of reward processing: from value representation, associative learning, and incentive salience to affective processing (Bartra et al., 2013; Berridge and Kringelbach, 2008; Chase et al., 2015).

Notably, the reward circuit is highly conserved in evolution, with remarkable similarities between humans and other mammals (Haber and Knutson, 2010; Loonen and Ivanova, 2016). Yet, the capacity to experience pleasure from aesthetic rewards, and particularly from music, is thought to be a unique human trait. Indeed, important differences exist between music and primary rewards (e.g. food). For instance, while hedonic reactions and preference for primary tastes, such as sweetness, are largely innate and highly preserved across species and individuals (Berridge, 2000), musical preferences are shaped by previous exposure, cognitive abilities, musical education, and cultural background; thus they are largely influenced by learning and plasticity (Gold et al., 2019b, 2019a; Greenberg et al., 2015; Haumann et al., 2018). In this regard, recent models hold that music-induced pleasure may be driven by anticipation and prediction mechanisms, which, in turn, have been linked to predictive coding theories (Cheung et al., 2019; Gold et al., 2019b; Koelsch et al., 2019; Salimpoor et al., 2015). In line with this idea, neuroimaging studies have shown that music-induced pleasure may be mediated by the crosstalk between the ancient reward circuitry and higher-order cortical regions involved in auditory cognition and predictive coding that are phylogenetically newer and especially well developed in humans, such as the superior temporal gyrus and the inferior frontal gyrus (Loui et al., 2017; Martínez-Molina et al., 2016; Sachs et al., 2016; Salimpoor et al., 2013).

The engagement of both higher-order neural mechanisms and the reward system appear to be a hallmark of musical reward as compared to primary and secondary rewards. However, due to the lack of comparative research comparing music and other types rewards it is unclear to what extend this is the case. Among all other reward types, hedonic responses to food represent a great control condition to assess common and distinct brain responses associated with music-induced pleasure since (i) the brain correlates of food reward have been extensively investigated in both humans and non-humans animals (particularly hedonic responses to sweetness) and (ii) food is passively delivered in real-time in human neuroimaging studies, as it is the case of music. Therefore, a direct comparison between these two reward types may help us to disentangle specific- and common-brain activations during their consumption. Indeed, a part from differences in the engagement of high-order cortical regions, differences between both may be present within the reward circuitry as well. For instance, although various parts of the striatum – the caudate, the putamen and the nucleus accumbens – have been shown to be activated by both music and food, it’s not clear whether one predominates over the other as function of the type of reward. Music reward studies have particularly emphasized the role of both the caudate and the nucleus accumbens in music-induced pleasure (Salimpoor et al., 2011), while food rewards might recruit more lateral portions of the striatum as well (Delgado, 2007). In addition, previous studies on primary and secondary rewards have shown a gradient along the posterior-anterior axis of the ventral prefrontal cortex as a function of the abstractness of the reward, with anterior parts – phylogenetically newer than posterior and medial parts – encoding more abstract representations (Klein-Flügge et al., 2013; O’Doherty et al., 2001; Rudebeck and Murray, 2014; Sescousse et al., 2013, 2010). Thus, one fundamental hypothesis is that music rewarding experiences would be represented in more anterior areas of the prefrontal cortex than food rewards.

In order to address these questions we directly compared brain activations to food and music reward to identify both overlapping and specific reward-type activations by using a coordinate-based meta-analytic approach. Specifically, we conducted a meta-analysis of neuroimaging studies that investigated brain responses to either music- or food-induced pleasure. In order to identify hedonic-related activations, all studies included in the meta-analysis involved either music or food “consumption” inside the scanner, and reported contrasts exploring brain activations as a function of subjective reports of pleasure. By using this approach, we focus directly on hedonic or affective processes rather than perceptual or cognitive aspects of music vs food. We first assessed the brain regions that were consistently engaged by either music or food reward, and next, we investigated common and specific reward type activations. We hypothesized that while food and music-induced pleasure would show some degree of overlapping activations within the reward network, music-induced pleasure would involve the engagement of higher-order cortical regions; we also hypothesized that differences may be present in the location and the predominance of one reward-type over the other within the reward circuitry, particularly in the ventral prefrontal cortex and the striatum (Delgado, 2007; Sescousse et al., 2010).

## METHODS

### Study selection for the meta-analysis

We conducted two independent searches (one for food studies, another for music studies) in PubMed in order to identify fMRI and Positron Emission Tomography (PET) studies exploring music and food-induced pleasure. Our search included a combination of the following: (i) the type of reward [i.e. “music”; (“food” OR “taste” OR “juice”)]; (ii) reward-related terms [i.e. (“reward” OR “pleasure” OR “liking”)]; and (iii) neuroimaging terms [i.e. (“fMRI” OR “PET” OR “neuroimaging” OR “cerebral blood flow”)]. The search on food reward was performed from 2012 to present. Studies on food reward prior to 2012 were obtained from a previous meta-analysis on food reward (Sescousse et al., 2013). The searches led to 87 studies on music and 532 on food reward. Articles were first selected by their title and abstract, and then carefully read to make sure that they fulfilled the following selection criteria:

1. Whole-brain results were reported (excluding studies entirely based on ROI analyses, without full-brain coverage, and PET studies using selective radiotracers other than H215O).
2. Involved healthy, drug-free, adult subjects (excluding studies performed on children, teenagers and old adults and studies with fewer than 10 subjects).
3. Involved the delivery, in real time, of either food/tastes or music.
4. Participants reported subjective ratings of pleasure, liking or preference.
5. Contrasts reported reflected hedonic processing at the time of music listening or food consumption (including contrasts such as like>dislike, positive correlations with pleasantness, correlation with individual differences in hedonic processing). Contrasts based on baseline subtraction were not included to avoid nonspecific effects.

Seventeen experiments (190 foci/330 subjects) on music rewards and twenty-one experiments (193 foci/373 subjects) on food reward (Tables 1–2) fulfilled our inclusion criteria.

**Table 1.** Music reward studies included in the meta-analysis.

| Study | Modality | n | Foci | Task | Music selection | Contrast |
| --- | --- | --- | --- | --- | --- | --- |
| Blood et al., 1999 | PET | 10 | 2 | Listening to 6 versions of a novel musical passage varying in degree of dissonance | Experimenter | Positive correlation with pleasantness |
| Blood et al., 2001 | PET | 10 | 10 | Listening to music that elicits chills | Participants | Positive correlation with chill intensity |
| Brattico et al., 2016 | fMRI | 29 | 21 | Listening to liked and disliked musical pieces with happy and sad connotation | Participants | Like > Dislike |
| Caria et al., 2011 | fMRI | 14 | 20 | Listening to sad and happy music | Participants & Experimenter | Favorite > Standard (Control group) |
| Gold et al., 2019 | fMRI | 19 | 11 | Decision-making task leading to consonant and dissonant musical endings | Experimenter | Consonance > Dissonance |
| Huang et al., 2016 | fMRI | 18 | 26 | Listening to artistic and popular music, and musical notes | Experimenter | Popular music > Notes clip<br>Artistic music > Notes clip |
| Keller et al., 2013 | fMRI | 21 | 12 | Listening to both familiar and unfamiliar music, and scrambled versions | Experimenter | Negative correlation with anhedonia trait |
| Kim et al., 2017 | fMRI | 23 | 7 | Listening to consonant and dissonant music | Experimenter | Consonant > Dissonant |
| Koelsch et al., 2006 | fMRI | 11 | 11 | Listening to pleasant music and pitch-shifted versions | Experimenter | Pleasant > Unpleasant |
| Martinez-Molina et al., 2016 | fMRI | 29 | 13 | Listening to both pleasant and unpleasant music while reporting real-time ratings of pleasure | Participants & Experimenter | Positive correlation with pleasantness (excluding musical anhedonics) |
| Mas-Herrero et al., *in prep* | fMRI | 17 | 10 | Listening to pleasant music while reporting real-time ratings of pleasure | Participants & Experimenter | Positive correlation with pleasantness at sham |
| Montag et al., 2011 | fMRI | 33 | 6 | Listening to subject-selected favorite and unlike songs | Participants | Pleasant > Unpleasant |
| Mueller et al., 2015 | fMRI | 23 | 16 | Listening to original (pleasant) and manipulated (unpleasant) music | Experimenter | Pleasant > Unpleasant |
| Pereira et al., 2011 | fMRI | 14 | 8 | Listening to a list of pop/rock music | Experimenter | Like > Dislike |
| Salimpoor et al., 2013 | fMRI | 19 | 8 | Listening to new released music with the possibility to buy the actual music | Experimenter | Positive correlation with the amount of money willing to pay |
| Trost et al., 2014 | fMRI | 20 | 7 | Listening to either consonant or dissonant piano music while performing a speeded visuo-motor detection task | Experimenter | Consonant > Dissonant |
| Wallmark et al., 2018 | fMRI | 20 | 2 | Listening to both familiar and unfamiliar music, either pleasant or unpleasant. | Experimenter | Like > Dislike |

**Table 2.** Food reward studies included in the meta-analysis.

| Study | Modality | n | Foci | Task | Contrast |
| --- | --- | --- | --- | --- | --- |
| Berns et al., 2001 | fMRI | 25 | 1 | Passive delivery task | Preferred > Nonpreferred |
| de Araujo et al., 2003 | fMRI | 11 | 8 | Passive delivery task | Sucrose > Tasteless drink |
| Felsted et al., 2010 | fMRI | 40 | 13 | Passive delivery task | Milkshake > Tasteless drink |
| Grabenhorst et al., 2010 | fMRI | 14 | 6 | Passive delivery task | Positive correlation with pleasantness of milkshake flavour |
| Grabenhorst et al., 2010b | fMRI | 12 | 3 | Passive delivery task | Positive correlation with pleasantness |
| Green and Murphy, 2012 | fMRI | 12 | 24 | Passive delivery task | Sucrose > Water (soda drinkers) |
| Green and Murphy, 2012 | fMRI | 12 | 15 | Passive delivery task | Sucrose > Water (non-soda drinkers) |
| Haase et al., 2009 | fMRI | 18 | 36 | Passive delivery task | Sucrose > Water |
| Kim et al., 2011 | fMRI | 18 | 4 | Instrumental task | Positive correlation with the value of juice |
| Kishi et al., 2017 | fMRI | 21 | 6 | Passive delivery task | Sweet solution > Water |
| McCabe & Rolls, 2007 | fMRI | 12 | 2 | Passive delivery task | Positive correlation with taste pleasantness |
| McCabe et al., 2011 | fMRI | 15 | 9 | Passive delivery task | Chocolate > Tasteless drink |
| McCabe et al., 2013 | fMRI | 16 | 12 | Passive delivery task | Chocolate > Tasteless solution |
| Monteleone et al., 2017 | fMRI | 20 | 13 | Passive delivery task | Sucrose > Water |
| Murray et al., 2014 | fMRI | 20 | 5 | Passive delivery task | Chocolate > Control |
| Nolan-Poupert et al., 2013 | fMRI | 20 | 3 | Passive delivery task | Milkshake > Tasteless solution |
|  |  |  | 3 | Passive delivery task | Positive Correlation with ad Libitum Intake |
| Rolls & McCabe, 2007 | fMRI | 16 | 6 | Passive delivery task | Chocolate > Tasteless drink |
| Small et al., 2008 | fMRI | 12 | 4 | Passive delivery task | Juice > Tasteless drink |
| Spetter et al., 2012 | fMRI | 15 | 3 | Passive delivery task | Sweet & Savory > water |
| van den Bosch et al., 2014 | fMRI | 34 | 14 | Passive delivery task | Sucrose > Water |
| Zald et al., 1998 | PET | 10 | 3 | Passive delivery task | Chocolate > Water |

### Meta-analysis methods

We conducted two independent coordinate-based meta-analyses, one for music experiments and one for food experiments, using anisotropic effect-size Seed-Based *d* Mapping (SDM, version 5.15, formerly “Signed Differential Mapping”) meta-analysis (http://www.sdmproject.com). Using peak coordinates, SDM generates a voxel-level map of effect sizes by converting the reported *t* values to effect size (Hedge’s *d*) and modeling an anisotropic kernel. All individual effect size maps are then combined with a random-effects model, accounting for sample size and effect size variability within and between studies. SDM *Z* maps are then recreated dividing meta-analytic effect sizes by their standard errors across studies. Since the resulting *Z* values do not typically follow a normal distribution, SDM uses permutation statistics to estimate a null distribution. All the analyses conducted were based on 50 permutations. Finally, statistical thresholding was performed using voxel-level *p*< 0.001 uncorrected, peak SDM-Z > 1, and a minimum of 10 continuous voxels.

Primary analyses examined brain activity consistently engaged in either studies of music- or food-induced reward. The two resulting maps were further binarized and combined to identify “overlapping” brain regions. Next, we formally tested whole-brain differences between both reward-types by calculating the difference between both rewards in each voxel and determining its statistical significance using a randomization test as implemented in SDM.

We additionally performed Jackknife sensitivity analyses to examine the robustness of the results and conducted Egger tests on each cluster to assess potential publication bias (Forero et al., 2019).

## RESULTS

We first examined brain regions that were consistently engaged when listeners reported a pleasurable experience to music. As illustrated in Figure 1, the analysis yielded a set of brain regions including the bilateral insula (INS), the bilateral superior temporal gyrus (STG), the right inferior frontal gyrus (IFG), the bilateral ventral striatum, the anterior prefrontal cortex, and the ventro-medial prefrontal cortex (vmPFC, whole-brain maps of the main results are available online at Neurovault: https://identifiers.org/neurovault.collection:8567).

**Figure 1.**
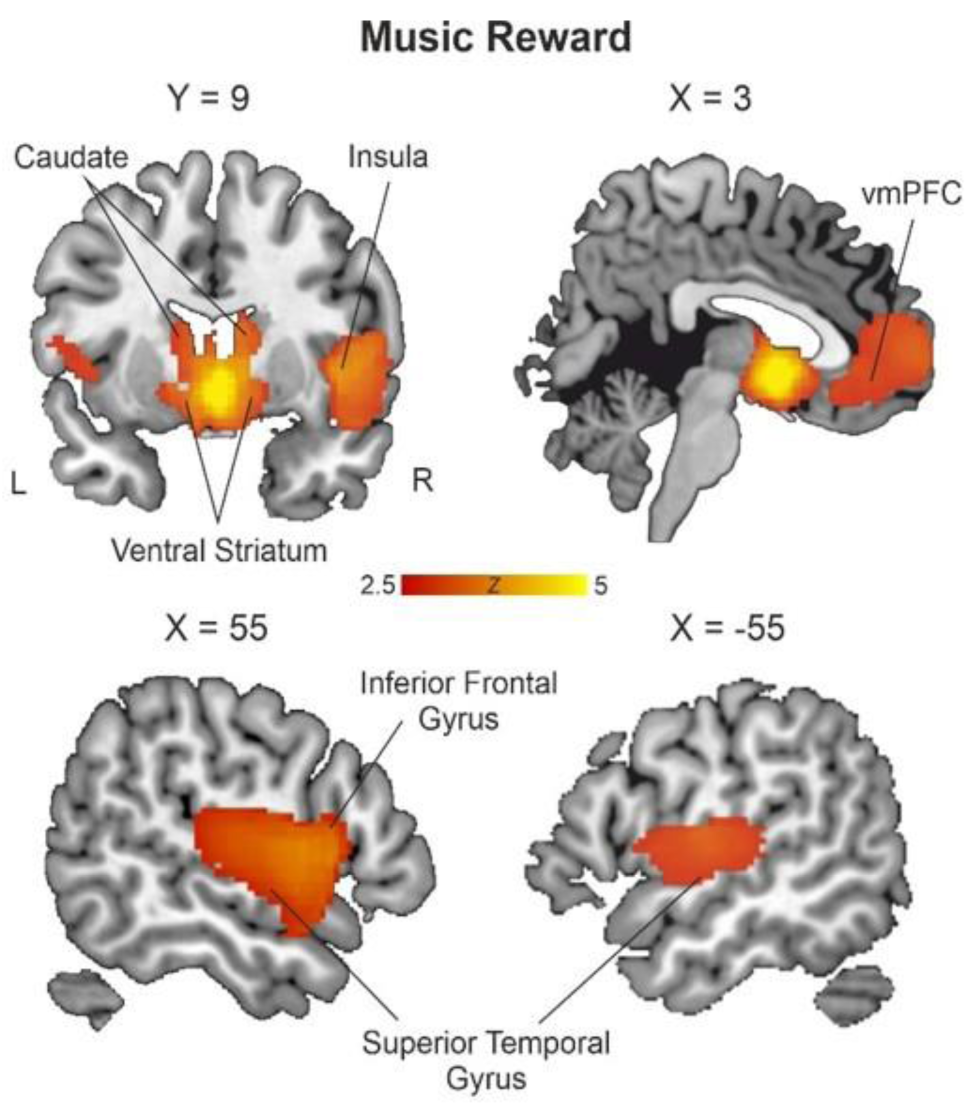
Whole-brain meta-analysis of studies investigating music-induced pleasure

Figure 2 shows the main findings for the meta-analysis of neuroimaging studies investigating food-induced pleasure. The analysis revealed engagement of the bilateral insula, putamen, amygdala, ventral striatum, somatosensory cortex, posterior cingulate cortex, vmPFC and thalamus.

**Figure 2.**
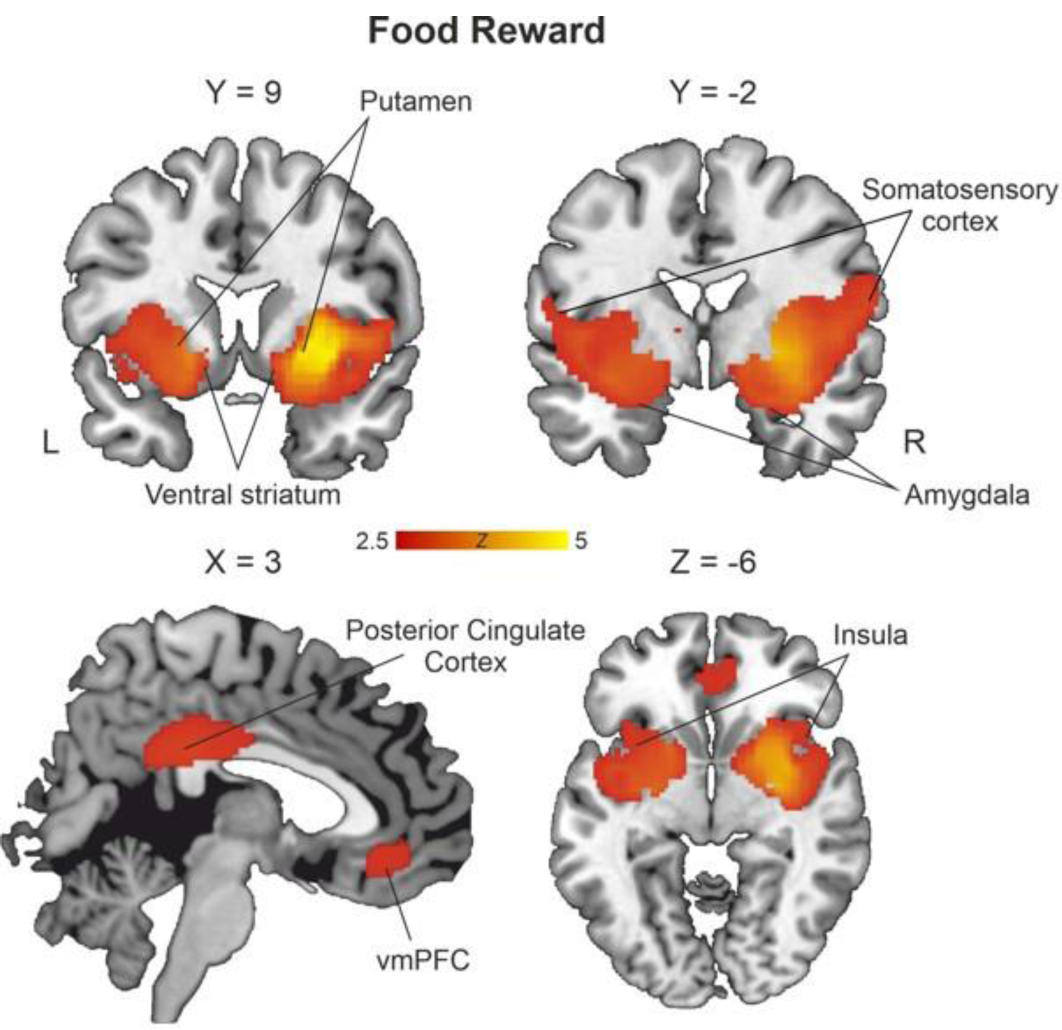
Whole-brain meta-analysis of studies investigating food-induced pleasure.

Using a jackknife procedure, we found that the two activation maps (food and music) were replicated in all jackknife analyses, reflecting the consistency and robustness of our results (Figure S1 and S2). In addition, funnel plots and Egger test in the clusters identified showed no evidence of publication bias for either music or food studies in any of the clusters identified (Figure S4 and S5).

Finally, to ensure that the effects in the music meta-analysis were not driven by those studies in which actions or decisions were made (instead of passively listening to music), we reran our analyses excluding the 5 experiments involving real-time ratings of pleasure, decision-making or action selection. The results remained qualitatively similar, suggesting that our main findings are not influenced by the active judgment and performance in these data sets (Figure S3).

Next, in order to identify those brain regions shared between both types of reward, we performed a conjunction between the two maps previously described (Figure 3). The analysis indicated that a set of brain regions were recruited in response to either food or music reward. These regions included the bilateral ventral striatum, the bilateral insula and the vmPFC.

Finally, in order to identify reward type-specific activations, we performed statistical comparisons between studies. The results showed that the right ventromedial striatum, the right STG and the anterior prefrontal cortex elicited a more reliable activation to music-induced pleasure compared with food. On the contrary, the left insula, bilateral putamen and right amygdala were more likely activated by food-induced pleasure than music.

## DISCUSSION

In the current study we performed a series of meta-analyses to examine overlapping and specific neural responses associated with music- and food-induced pleasure. This is the first attempt to explore brain regions consistently engaged in response to music-induced pleasure and to investigate partial dissociations with respect to food rewards. Previous meta-analyses on music (Koelsch, 2014) have mixed together contrasts reflecting pleasantness, unpleasantness, and music-induced sadness and happiness, among others, as well as different neuroimaging modalities (PET imaging of dopamine-D2 receptor, functional connectivity and fMRI activations). Here, we focused on hedonic-related fMRI activations in order to specifically identify those brain regions that are more consistently engaged when individuals are experiencing pleasure induced by either music or food. Our findings are consistent with the existence of a common reward circuit, involving the ventral striatum, the insula, and the vmPFC. Yet, comparative analyses indicated a partial dissociation in the engagement of the reward circuitry as a function of the type of reward, as well as additional reward type-specific activations in brain regions related to perception, sensory processing, and learning.

### The striatum

Both food and music-induced pleasure were associated with activations of the bilateral ventral striatum (VS). Although its precise functional role is still a matter of fervid debate, a wealth of fMRI studies has identified the VS in reinforcement learning, reward valuation and incentive motivation, using either primary or secondary rewards (Bartra et al., 2013; Berridge, 2007; Chase et al., 2015; O’Doherty et al., 2004). Indeed, the VS is considered a key hub of the reward circuitry, with extensive connections to prefrontal regions and limbic structures (Haber and Knutson, 2010) and is one of the main inputs of the mesolimbic dopaminergic neurons.

The key role of the ventral striatum and the dopaminergic system in generating feelings of musical pleasure has been long emphasized. Concretely, previous studies have shown that (a) the degree of dopaminergic neurotransmission is related to the degree of music-induced pleasure (Salimpoor et al., 2011), (b) TMS-induced modulation of activity in the ventral striatum and its connectivity changes subjective feelings of pleasure bidirectionally (Mas-Herrero et al., 2018a, Mas-Herrero et al., *subm*), and (3) pharmacological dopaminergic manipulations effectively induce changes in subjective pleasure, affecting the frequency and the intensity of music-induced chills (Ferreri et al., 2019).

Interestingly, hedonic-related activations in the ventral striatum were more consistent in studies involving music. Previous meta-analysis and studies comparing brain activations to primary (e.g. food) and secondary (e.g. money) rewards have reported similar differences, with the latter showing more robust activations in the ventral striatum (Bartra et al., 2013; Sescousse et al., 2013). This effect has been attributed to the fact that monetary rewards are generally delivered in instrumental contexts in which the ventral striatum may preferentially encode reward prediction errors signals conveyed by dopaminergic neurons (Chase et al., 2015; Mas-Herrero et al., 2019; Schultz et al., 1997). In line with this idea, when food or erotic rewards are presented in an instrumental context, they tend to recruit the ventral striatum to a similar extent as monetary rewards (D’Ardenne et al., 2008; McClure et al., 2003; Sescousse et al., 2010). Yet, no instrumental context was present in most of the studies included in the music meta-analysis (15 out of 17). Even when studies involved action-selection, learning, and real-time ratings of pleasantness were excluded (n=5), a similar pattern of activations was found in response to music-induced pleasure (see Figure S3).

Thus, in the absence of any action or instrumental learning task, hedonic-related activations in the ventral striatum while passively listening to music were more consistent than in response to food reward. However, the presence of regularities in music may turn it into a learning scenario by itself, although not explicitly. Indeed, current neuro-functional models of musical pleasure suggest that the engagement of the ventral striatum, and the NAcc in particular, may be driven by music-induced expectations, particularly when those are violated (Gold et al., 2019a; Koelsch et al., 2019; Salimpoor et al., 2015; Shany et al., 2019). Notably, bodily reactions such as ‘chills’ which are generally associated with particularly intense and pleasurable responses to music, and have been associated with greater engagement of the NAcc (Grewe et al., 2009, 2005; Mas-Herrero et al., 2018b; Salimpoor et al., 2009), are often experienced following musical surprises (Grewe et al., 2007; Guhn et al., 2007; Harrison and Loui, 2014; Nagel et al., 2008; Panksepp, 1995; Sloboda, 1992). Finally, recent studies using an information-theoretic model of auditory expectation indicate that listeners may prefer music containing surprises, yet in a context in which those can be learned and anticipated (Cheung et al., 2019; Gold et al., 2019b). These findings resonate with learning theories that suggest that the information gain of balancing uncertainty and surprise (Oudeyer et al., 2016) could be intrinsically rewarded by the brain in order to foster our fundamental need of generating accurate models of the world. Thus, the information gain triggered by musical surprises and uncertainty, and our ability to successfully contextualize and resolve them, could be a potential underlying mechanism behind the rewarding properties of music and the consequent engagement of the reward circuitry in general and the VS in particular. In contrast to these aspects of music, which depend on the temporal unfolding of events over time and their anticipation, in the case of food reward there is no temporal sequencing of events, at least not in the way the studies are conducted. A food reward is presented but there are no cues that predict it nor is there any surprise involved.

Furthermore, food-related studies showed more consistent activations in the bilateral putamen, which receive important projections from somatosensory regions (Sgambato-Faure et al., 2016). Interestingly, food intake activates both taste and nutrition-sensing pathways. Recent models suggest that these pathways may be mediated by separate striatal circuits: while ventral regions of the striatum may encode the hedonic value of food, dorso-lateral regions may convey nutritional values reflecting the energy present in aliments (de Araujo, 2016; Tellez et al., 2016). On the other hand, studies performed on non-human primates comparing food and sex-motivated behavior have shown that loss of food motivation may be driven by alterations in the lateral portions of the VS, in contrast to sexual manifestations (erection) which may be driven by the functioning of the medial regions of the VS (Sgambato-Faure et al., 2016). This dissociation within VS has been related to the existence of separate fronto-striatal circuits associated to different motivation domains. The medio-lateral striatal dissociation between music and food reward found in our meta-analysis further supports the presence of distinct pathways associated with different rewards, and the relevance of specific-reward type circuits in motivated behaviors.

### Auditory cortical regions

The greater engagement of brain structures particularly involved in auditory cognition and predictive coding while listening to pleasant music, such as the right STG and the right IFG, further reinforces the idea that learning is crucial for the experience of musical pleasure. The right STG has been consistently implicated in various processes relevant for music perception, including pitch representation (Coffey et al., 2016; Johnsrude et al., 2000), tonal pattern processing (Foster and Zatorre, 2010a; Patterson et al., 2002), tonal working memory (Albouy et al., 2019), tonal learning (Herholz et al., 2016), and musical imagery (Herholz et al., 2012). Recent findings indicate that this right hemisphere specialization in music may arise from differential sensitivity to acoustical cues between the left and the right auditory cortices, with a specific sensitivity to spectral information in the right hemisphere and greater sensitivity to temporal information in the left (Albouy et al., 2020). Notably, our meta-analysis also indicates the engagement of the right IFG while listening to pleasant music, which goes in line with the right-hemispheric dominance in music processing. Previous fMRI and MEG studies have implicated the right IFG in musical structure processing – responding to musically unexpected events (Koelsch et al., 2005; Tillmann et al., 2006, 2003)–, and tonal working memory -while retaining, manipulating and retrieving tonal information (Albouy et al., 2018, 2017; Foster and Zatorre, 2010b). Consistent with the importance of these structures for music processing, individuals suffering from amusia (a deficit in music perception and production abilities, particularly in pitch processing) present morphological brain anomalies both in white and grey matter concentrations in the right IFG (Albouy et al., 2013) and in the connectivity between the STG and the IFG (Loui et al., 2009). It is important to note that the engagement of both the right STG and IFG in the current meta-analysis do not merely reflect perceptual or cognitive aspects related to general music processing (independently of its pleasantness) since all the contrasts included in the meta-analysis focus directly on hedonic or affective processes.

Previous fMRI studies have already shown that these high-order cortical regions involved in auditory cognition and predictive coding show enhanced coupling with reward-related structures, such as the NAcc, while listening to pleasant music (Salimpoor et al., 2013), coinciding with music-induced surprises (Shany et al., 2019). In addition, individual differences in music reward sensitivity are accompanied by similar differences in functional and structural connectivity between the right STG and reward- and emotion-related structures such as the vmPFC, the ventral striatum and the insula (Loui et al., 2017; Martínez-Molina et al., 2019, 2016; Sachs et al., 2016). In particular, specific musical anhedonics – individuals who do not derive pleasure from music but show intact music perceptual abilities as well as intact affective reactions to other reward types (Mas-Herrero et al., 2018b, 2014) – showed disrupted auditory-striatal coupling while listening to music. Conversely, individuals with high sensitivity to music presented the reverse pattern, with greater functional connectivity between auditory cortices and reward centers (Martínez-Molina et al., 2019, 2016). Finally, by combining fMRI with Transcranial Magnetic Stimulation (TMS) over the left dorso-lateral prefrontal cortex, a procedure known to modulate striatal function (Strafella et al., 2001), we have recently provided further evidence in favor of the causal role of auditory-striatal interactions in music-induced pleasure (Mas-Herrero et al., 2018a, Mas-Herrero et al., *subm.*). Specifically, TMS-induced changes in subjective musical pleasure were accompanied by modulation in the cross-talk between the right STG and the NAcc coinciding with the peak experience of musical pleasure.

Altogether, these findings fit well with the idea that the exchange of information between high-order cortical regions involved in auditory perception and cognition, on the one hand, and reward systems, on the other hand, plays a key role in musical pleasure.

### The ventro-medial Prefrontal Cortex

In a similar fashion as the ventral striatum, the vmPFC also responded to both food and music-induced pleasure. The ventral prefrontal cortex is a multisensory hub that receives inputs from sensory modalities such as taste, olfaction, audition, vision and somatic sensation and projects primarily to the ventral striatum (Haber and Knutson, 2010). Early fMRI studies have shown that vmPFC activations consistently correlates with subjective reports of pleasure in response to various types of rewards – including primary, secondary and abstract (Blood and Zatorre, 2001; Elliott et al., 2003; Knutson et al., 2001; O’Doherty et al., 2001; Rolls, 2000). The vmPFC is thought to integrate signals from different sensory modalities and represent them on a common scale, reflecting the attractiveness or value of rewards, for the purpose of comparison and evaluation (Kringelbach, 2005). In accordance with this view, both food and music-induced pleasure were associated with activations in the vmPFC (Fig 3). The cluster identified nicely overlaps with previous meta-analyses on subjective value comparing primary and monetary rewards, which also agrees with its role in representing reward value independently of the nature of the reward (Bartra et al., 2013; Sescousse et al., 2013).

**Figure 3.**
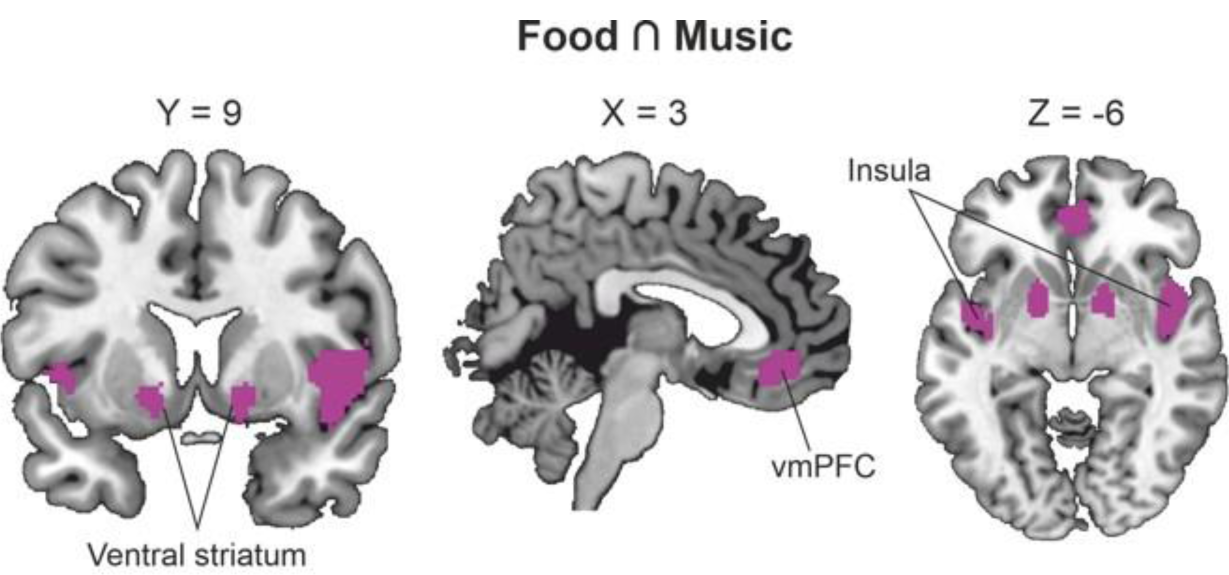
Overlap between both food- and music-induced pleasures.

In addition, our results show that music recruits anterior portions of the prefrontal cortex more reliably than food rewards (Fig 4). The cluster identified falls within the ventral portions of Broadmann area 10. Previous studies have implicated this region in the coordination of information processing in tasks involving multiple cognitive operations (Ramnani and Owen, 2004). Indeed, music requires the coordination and integration of several cognitive operations involving temporal expectations, predictions, working memory and learning across multiple hierarchical timescales. Notably, this area has considerable connections with the superior temporal gyrus (Petrides and Pandya, 2007; Saleem et al., 2008). Evidence in favor of the role of fronto-temporal connections in music-induced pleasure comes from patients with fronto-temporal lobe dementia (FTLD). Previous studies have shown that patients with FTLD may develop musicophilia, that is, a specific craving for music (Fletcher et al., 2015, 2013). This particular phenotype has been associated with changes in grey matter volume in a distributed network including the right temporal cortex and the anterior prefrontal cortex, among others (Fletcher et al., 2015, 2013). Overall, these findings together with our results suggest a key role of fronto-temporal pathways in musical pleasure.

**Figure 4.**
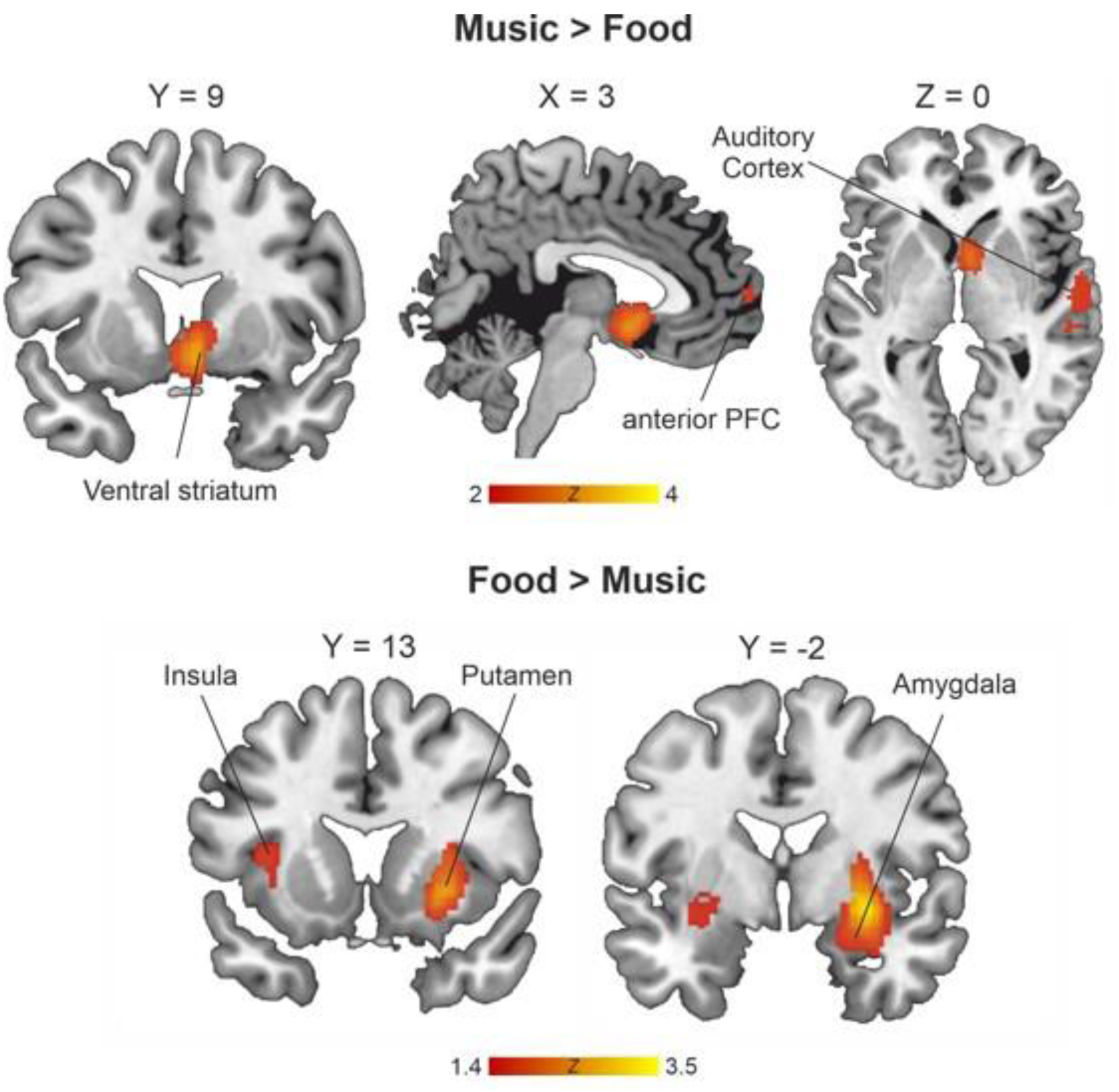
Brain regions more reliably activated by one reward compared to the other.

### The insula

The third brain region that was consistently engaged across both music and food rewards is the posterior insula bilaterally. The insular cortex also represents an integration hub and is at the crossroad of several circuits involved in sensory, cognitive, motivational and emotional functions (Benarroch, 2019; Gogolla, 2017; Uddin et al., 2017). Studies conducted in humans and non-human primates have identified a posterior-to-anterior connectivity gradient, with the anterior parts mostly connected to the frontal cortex and limbic regions, and the posterior portions heavily connected with sensorimotor, visual and auditory cortices (Benarroch, 2019; Gogolla, 2017; Uddin et al., 2017). These differences in connectivity pattern have been associated with different functionality: while anterior regions have been related to introspective awareness of emotion and bodily states (Critchley, 2004; Paulus and Stein, 2006), posterior sections have been involved in somato-sensory processing and integration (Rodgers et al., 2008; Stephani et al., 2011). In particular, previous studies have shown the pINS is activated by sounds (reflecting auditory responses that resemble those described in the auditory cortex, Blenkmann et al., 2019) and mouth movements (Woolnough et al., 2019). Thus, given the role of the pINS in somato-sensory function, the recruitment of the pINS by both food- and music-induced pleasure in our data may be reflecting greater auditory and sensorimotor processing to pleasant music and food, respectively. On the other hand, our results indicate that the anterior insula (aINS) was robustly and specifically activated by food rewards. The aINS also includes the primary taste cortex which receive multiple sensory inputs of gustatory cues, such as smell, taste, and texture, among others (de Araujo and Simon, 2009). This finding is consistent with the activations found in the somatosensory cortex in the meta-analysis of neuroimaging studies investigating food-induced pleasure (Figure 2). Similar to models of music, food-induced reward may rely on the interaction between brain regions involved in food perceptual processing and the common reward circuitry. Altogether, the results highlight the existence of specific routes of access into the reward circuitry depending on the modality and nature of the stimulus, which may provide a plausible mechanism explaining the existence of specific anhedonias (and hyperhedonias) in response to a particular type of reward.

### The amygdala

Our meta-analysis also revealed food-specific activations in the amygdala. The amydgala is especially well-known to mediate emotional processing, particularly aversive emotional reactions such as fear (LeDoux, 2000). Yet, accumulating evidence indicates that the amygdala is equally sensitive to rewarding stimuli (Bermudez and Schultz, 2010; Janak and Tye, 2015; Sugase-Miyamoto and Richmond, 2005). In addition, the engagement of the amygdala seems to scale with the emotional intensity and salience of the stimuli, regardless of their valence (Bonnet et al., 2015; Lang and Bradley, 2013). These findings have led to the idea that the predominant role of the amygdala may be the detection of motivationally salient information, rather than a specific role in valence processing as it was initially thought. Particularly, the amygdala may be involved in assigning an emotional tag to salient stimuli in order to guide decision-making (Gottfried et al., 2003), and acquire and retain lasting memories involving emotional experiences (Inman et al., 2018). Notably, we did not find amygdala activation associated with music-induced pleasure. This finding may be considered surprising given the rich emotional content of music. Yet, one potential explanation is that this lack of effect may be driven by differences in the fMRI contrasts generally used in music studies as compared to those employed in food-reward studies. While most of the food studies included in our meta-analysis compared brain activations between pleasant (sugar, chocolate, juice, etc) and neutral food (tasteless solutions, water, etc); most of the fMRI studies in music-induced pleasure compared pleasant to unpleasant music (particularly dissonant music). In this regard, dissonant and unpleasant music has been shown to activate paralimbic structures, including the amygdala as well as the parahippocampal gyrus (Blood et al., 1999; Gosselin et al., 2007; Koelsch et al., 2006a). Therefore, the dissociation found in the amygdala between food- and music-induced pleasures may reflect an imbalance in emotional salience between pleasant and neutral food and an equivalent emotional intensity between pleasant and unpleasant music. In addition, in most food studies, individuals were tested after a short period of fasting, which likely contributed to experiencing the actual food as more emotionally arousing. Therefore, the amygdala activations that we found in response to food-induced pleasure are more likely to reflect discrepancies in the emotional impact between target and baseline conditions in food-reward studies as compared with music-reward studies, rather than a specific role in the hedonic evaluation of primary rewards.

## Conclusions

The current meta-analysis is the first to systematically compare the neuroanatomical substrates of both a primary reward (food) and an abstract aesthetic reward (music). Our results indicate the existence of a set of brain regions that are consistently engaged while either listening to pleasant music or savoring palatable food. These brain regions constitute a common reward circuitry and include the bilateral striatum, the ventro-medial prefrontal cortex and the bilateral insula. However, comparative analysis indicated that the location of the activity within these regions varied somewhat across both rewards and the presence of reward type-specific activations. Overall, the current results support the idea that hedonic reactions rely on the engagement of a common reward network, yet through specific routes of access depending on the modality and nature of the input. In addition, the current findings support the interpretation that music-induced pleasure relies on the engagement of both higher-order cortical regions involved in auditory cognition and predictive coding, and reward-related structures.

## Akcnowledgements

R.J.Z. is supported by funds from the Canadian Institutes of Health Research (CIHR), the Natural Sciences and Engineering Research Council of Canada, and the Canada Fund for Innovation. E.M.-H. is supported by fundsfrom the Institutode SaludCarlos III through the Sara Borrell call, grantnumber CD19/00093(Co-fundedbythe European Social Fund. “ESFinvestinginyour future”). The funders had no role in the conceptualization, design, data collection, analysis, decision to publish or preparation of the manuscript

