## Supplemental Information for "Common and distinct neural correlates of music and food-induced pleasure: a coordinate-based meta-analysis of neuroimaging studies"

### Supplementary Information

**Figure S1.** Binarised jackknife analyses for music-induced pleasure. Leave-one-out validation indicated that the clusters identified in our meta-analysis are observed in all jackknife analyses. In red, the brain regions overlapping in all jackknife analyses at a threshold of $p < .005$ and $k \geq 10$ (17 analyses in total).

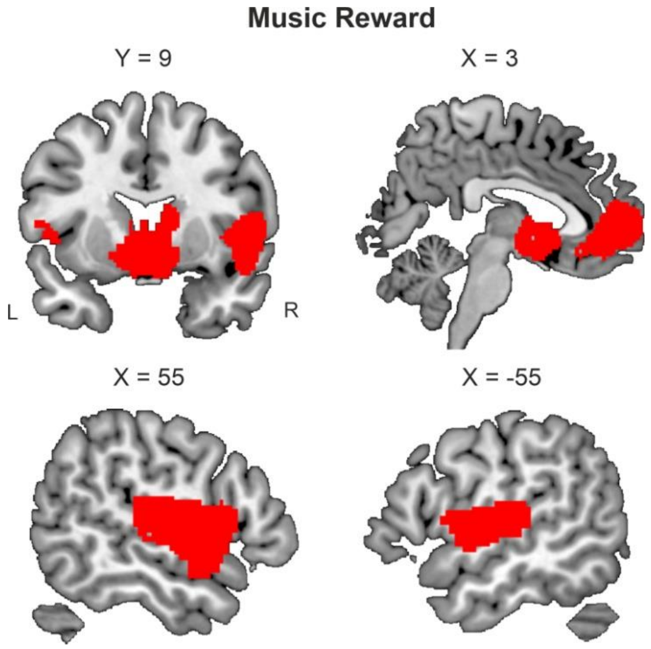

**Figure S2.** Binarised jackknife analyses for food-induced pleasure. Leave-one-out validation indicated that the clusters identified in our meta-analysis are observed in all jackknife analyses. In red, the brain regions overlapping in all jackknife analyses at a threshold of $p < .005$ and $k \geq 10$ (21 analyses in total).

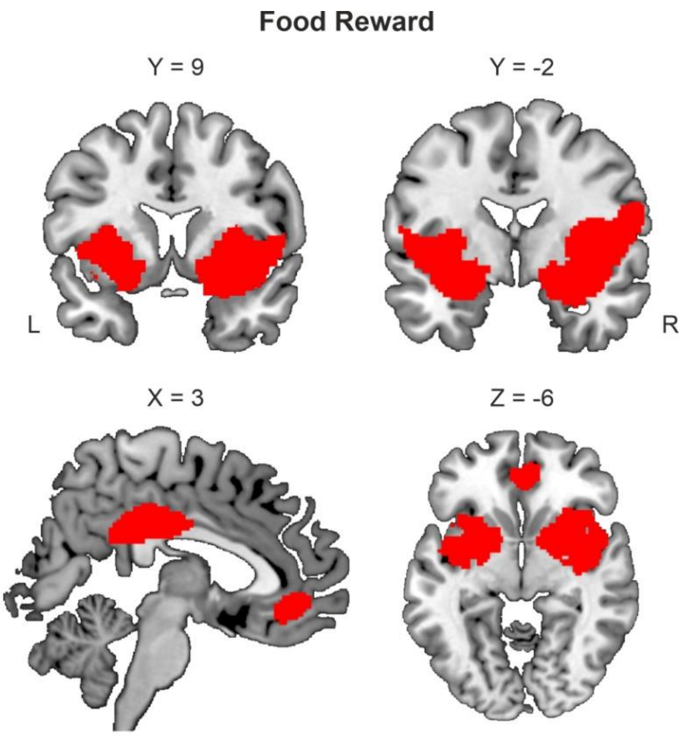

**Figure S3.** Stability of whole-brain meta-analytic results after excluding the 5 data-sets from the music-induced pleasure meta-analysis that explicitly involved learning, decision-making or action selection rather than passive listening. Note that the main findings are still present after the exclusion.

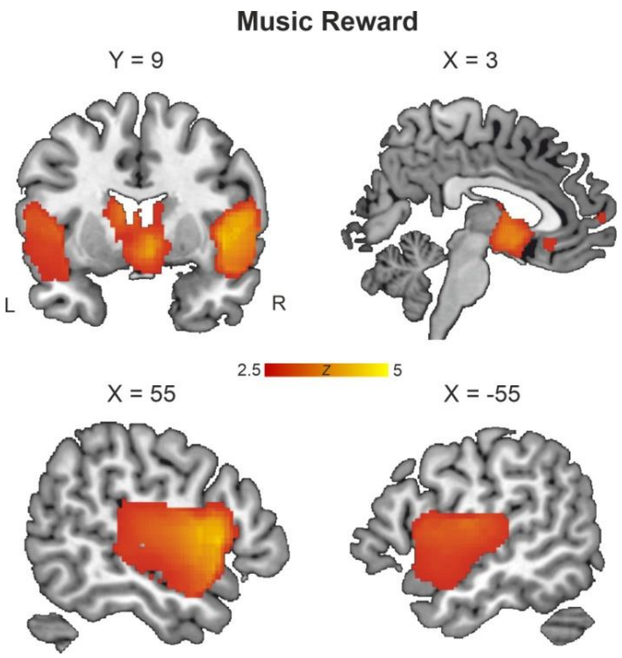

**Figure S4.** Funnel plots and Egger test assessing publication bias of fMRI studies investigating music-induced pleasure for each cluster identified.

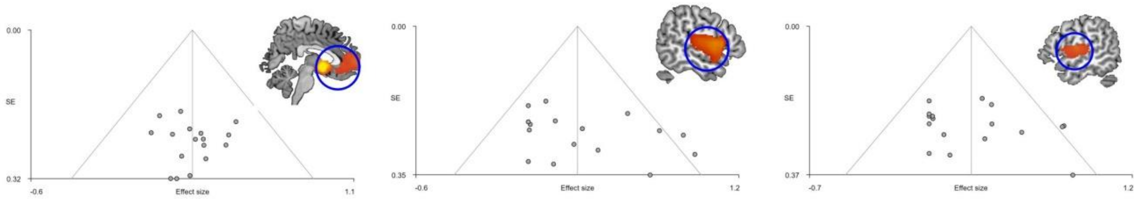


Bias: 0.17, t: 0.21, df: 15, p: 0.837

Bias: 2.60, t: 1.61, df: 15, p: 0.128

Bias: 1.66, t: 1.06, df: 15, p: 0.305

**Figure S5.** Funnel plots and Egger test assessing publication bias of fMRI studies investigating food-induced pleasure for each cluster identified.

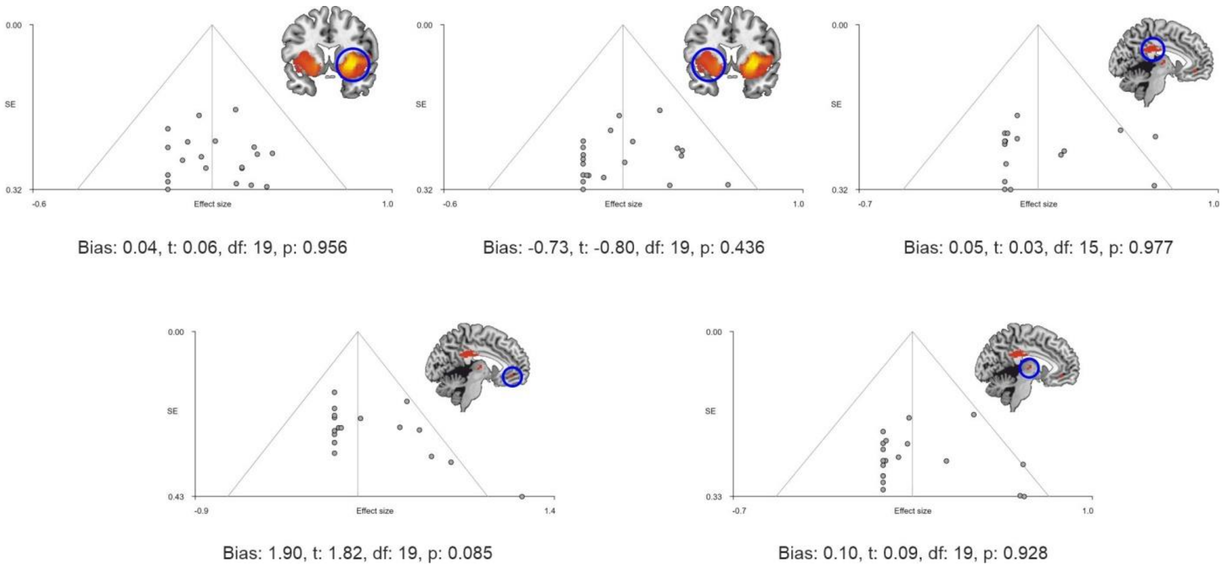
